# Automatically annotated motion tracking identifies a distinct social behavioral profile following chronic social defeat stress

**DOI:** 10.1101/2022.06.23.497350

**Authors:** Joeri Bordes, Lucas Miranda, Maya Reinhardt, Lea Maria Brix, Lotte van Doeselaar, Clara Engelhardt, Benno Pütz, Felix Agakov, Bertram Müller-Myhsok, Mathias V. Schmidt

**Affiliations:** Research Group Neurobiology of Stress Resilience, Max Planck Institute of Psychiatry, 80804 Munich, Germany; Research Group Statistical Genetics, Max Planck Institute of Psychiatry, 80804 Munich, Germany; International Max Planck Research School for Translational Psychiatry (IMPRS-TP), 80804 Munich, Germany; Pharmatics Limited, Edinburgh EH16 4UX, Scotland, UK

**Keywords:** chronic social defeat stress, machine learning, DeepOF

## Abstract

Severe stress exposure is a global problem with long-lasting negative behavioral and physiological consequences, increasing the risk of stress-related disorders such as major depressive disorder (MDD). An essential characteristic of MDD is the impairment of social functioning and lack of social motivation. Chronic social defeat stress is an established animal model for MDD research, which induces a cascade of physiological and social behavioral changes. The current developments of markerless pose estimation tools allow for more complex and socially relevant behavioral tests, but the application of these tools to social behavior remains to be explored. Here, we introduce the open-source tool “DeepOF” to investigate the individual and social behavioral profile in mice by providing supervised and unsupervised pipelines using DeepLabCut annotated pose estimation data. The supervised pipeline relies on pre-trained classifiers to detect defined traits for both single and dyadic animal behavior. Subsequently, the unsupervised pipeline explores the behavioral repertoire of the animals without label priming, which has the potential of pointing towards previously unrecognized motion motifs that are systematically different across conditions. We here provide evidence that the DeepOF supervised and unsupervised pipelines detect a distinct stress-induced social behavioral pattern, which was particularly observed at the beginning of a novel social encounter. The stress-induced social behavior shows a state of arousal that fades with time due to habituation. In addition, while the classical social avoidance task does identify the stress-induced social behavioral differences, both DeepOF behavioral pipelines provide a clearer and more detailed profile. DeepOF aims to facilitate reproducibility and unification of behavioral classification of social behavior by providing an open-source tool, which can significantly advance the study of rodent individual and social behavior, thereby enabling novel biological insights as well as drug development for psychiatric disorders.

## Introduction

Stress is an essential aspect of our daily lives, which contributes to our mood and motivation. However, exposure to severe stress can have negative consequences, which has become an increasing burden on society. In particular, stress-related disorders, such as major depressive disorder (MDD), have been steadily on the rise for the last decade (1). Our understanding of the behavioral and neurobiological mechanisms related to MDD is limited, which is a part of the reason for the moderate success of current drug treatments (2). MDD is a complex and heterogeneous disorder and the classification is dependent on a widespread set of symptoms. An important characteristic of MDD is the impairment of social functioning and lack of social motivation, which can lead to social withdrawal from society in extreme cases (3). In addition, disturbances in social behavior are an important risk factor for developing MDD, as poor social networks are linked to lowered mental and physical health (4, 5). The impact of social interactions was highlighted during the COVID-19 pandemic, where a substantial part of society experienced less to no social interactions for a sustained period of time. An increasing number of studies are now reporting the enormous impact of the pandemic, emphasizing a dramatic increase in the prevalence of stress-related disorders and in particular for MDD (6, 7). Unfortunately, there is still a lack of awareness of the importance of social interactions and their role in stress-related disorders. Therefore, it is crucial to increase the understanding of the biological and psychological mechanisms behind MDD and in particular the influence of social behavior on the development of MDD. Animal models have an important role in MDD research; although it is not possible to recreate the exact disorder as in humans, they do provide a controlled environment where symptoms of MDD can be investigated (8). The well-established chronic social defeat stress (CSDS) paradigm is continuously used for studying symptoms of MDD in animals (9, 10). In the CSDS model, mice are subjected daily to severe physical and non-physical stressors from aggressive mice for several weeks, which results in the chronic activation of the physiological stress response system, leading to bodyweight differences, enlarged adrenals, and elevated levels of corticosterone (11). In addition, animals subjected to CSDS show stress-related behaviors such as social avoidance, anhedonia, reduced goal-directed motivation, and anxiety-like behavior (9, 12–14). In particular, the CSDS-induced social avoidance behavior, which is the avoidance of a novel conspecific, is a recognized phenomenon that is used to investigate the social neurobiological mechanisms related to chronic stress exposure and stress-related disorders (10, 15, 16). Currently, several social behavior tasks can assess different constructs of social behavior, but especially the social avoidance task is well-established (16). It is important that these behavioral tasks are conducted with control over the environment to investigate the effects of external stimuli, such as stress exposure. For decades there has been a trend to standardize and simplify these tests in order to allow for greater comparability and higher throughput. Unfortunately, this has led to an oversimplification of the social behavioral repertoire and increased the risk for cross-over effects by other types of behavior, such as anxiety-related behavior. Moreover, due to limitations in tracking software, the analysis of the interaction between multiple freely moving animals remained difficult, which further limited the complexity of the behavioral assessment. Social behavior is a complex behavioral construct, relying on many different types of behavioral interactions, which often are too complicated, time-intensive, and repetitive to manually assess (17–19). Ultimately, this can lead to poor reproducibility of the social behavioral construct, as observed for social approach behavior (20). The current advancement in automatically annotated behavioral assessment, however, allows for high-throughput analysis using pose estimation, involving both supervised classification (intending to extract pre-defined and characterized traits) and unsupervised clustering (which aims to explore the data and extract patterns without external information) (21–26). Importantly, the open-source tool “DeepLabCut” has provided a robust and easily accessible system for deep learning motion tracking and markerless pose estimation (27, 28). The use of supervised classification, by defining the types of behavior a priori, is a powerful tool that simplifies the analysis by using predefined relevant behavioral constructs without losing the complexity of social behavior. Furthermore, recent studies have shown the value of unsupervised clustering in addition to the supervised analysis pipeline, as it can deal with the discovery of novel traits in a more exploratory fashion, which could reveal novel and more complex structures of behaviors (17, 22, 29–31). In addition, both the supervised and unsupervised pipelines can provide more transparency for the behavioral definition and can easily be shared via online repositories, which contributes to a more streamlined definition of behavior across different labs (19, 32). In the current study, we provide an application of our open-source tool “DeepOF”, which enables users to delve into the individual and social behavioral profiles of mice using DeepLabCut annotated pose estimation data. We do so by fitting both supervised classifiers capable of recognizing a set of predefined traits and by embedding our motion data in an unsupervised behavioral space, capable of explaining differences between both conditions without any label priming. Furthermore, DeepOF can retrieve unsupervised clusters of behavior that can be compared across conditions and therefore can hint at previously unrecognized behavioral patterns that trigger new hypotheses. As a proof of principle, we describe a distinct social behavioral profile following CSDS in mice that can be recapitulated with both supervised and unsupervised workflows. Moreover, we observe a clear state of arousal in social interaction tasks that fades over time and provide tools for the quantification of optimal behavioral differences across time and experimental conditions.

## Methods

### Time series extraction from raw videos

Time series were extracted from videos using DeepLabCut version 2.2b7 (single animal mode). 14 body parts per animal were tagged, including the nose, left and right ears, three points along the spine (including the center of the animal), all four extremities, the tail base, and three equidistant points along the tail (Figure 1A). The DeepLabCut model was trained to track up to two animals at once (one CD1 mouse and one C57Bl/6N mouse) and can be found in the supplemental material. Furthermore, with the multi-animal DeepLabCut (28), extending the tracking to animals from the same species is also possible.

**Fig. 1.**
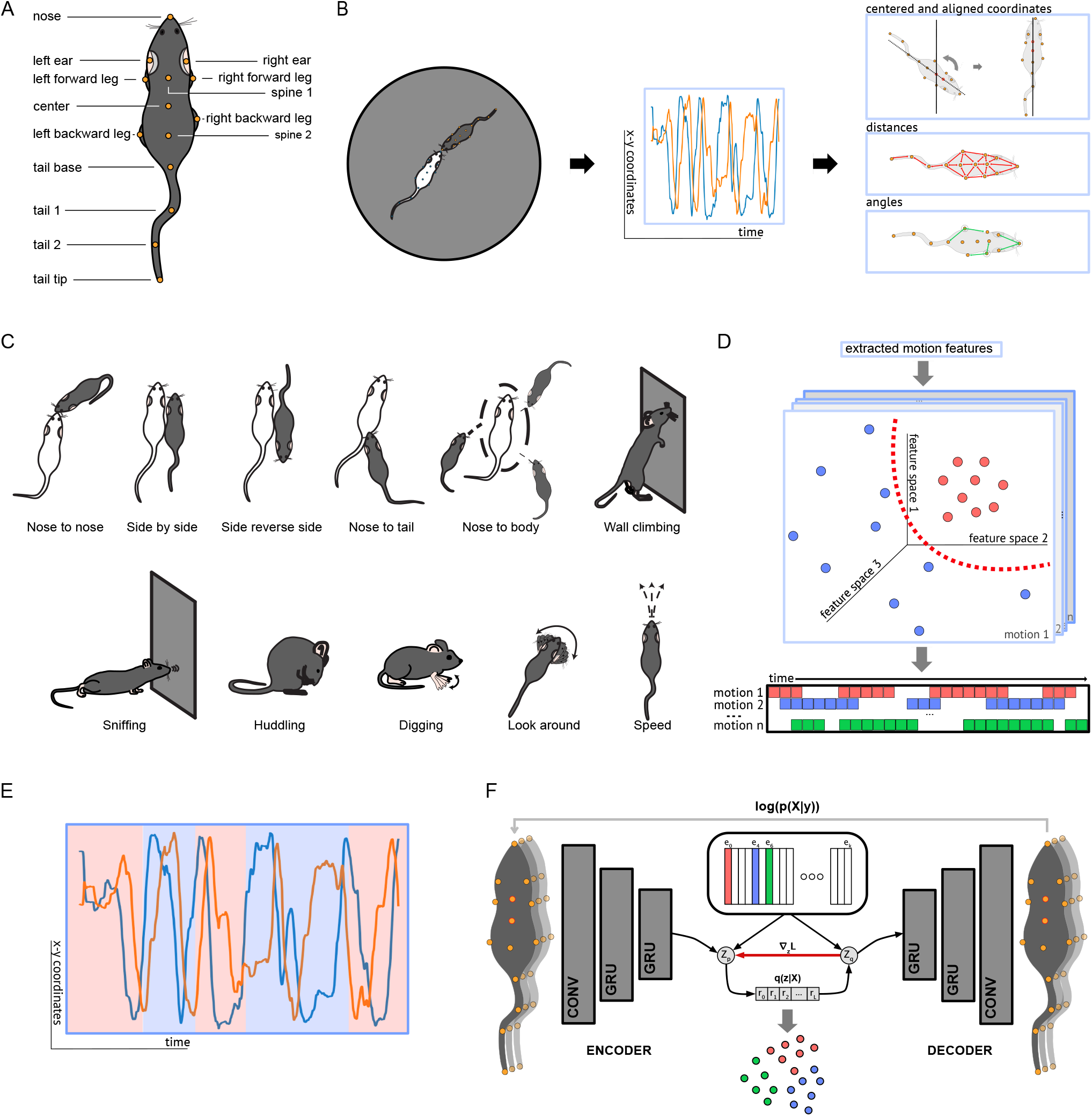
DeepOF workflow. A) 14 labels were tagged on each annotated mouse using DeepLabCut. B) The DeepOF preprocessing pipeline. Two mice (a C57Bl/6N experimental subject and a CD1 social companion) were tagged using the provided DeepLabCut models. Cartesian coordinates of all body parts were extracted using DeepLabCut. Next, DeepOF was used to smooth the retrieved trajectories, interpolate outliers, and extract a set of features, including centered and aligned coordinates, distances, and angles between body parts, as well as speed and acceleration. C) Set of motifs that the applied DeepOF supervised pipeline is able to retrieve. These include dyadic motifs (such as nose-to-nose and nose-to-body contacts) and individual motifs (climbing, digging, etc.), which are reported for all tracked mice individually. D) Schematic representation of the supervised pipeline in DeepOF. A set of extracted motion features (such as the coordinates, distances, and angles shown above; only 3 hypothetical dimensions are shown for visualization purposes) are fed to a set of pre-trained classifiers, which were obtained by following a thorough model selection pipeline on standardized data (see DeepOF documentation for details). Each of these classifiers reports the presence or absence of each behavioral trait at each time by learning how the corresponding trait is distributed in the feature space (red dots). The set of classifiers finally yields a table indicating the presence or absence of each motif across time, which can be used for further processing. Note that motifs are not necessarily mutually exclusive, as several predictors can be triggered at the same time. E) Illustration of the change-point detection pipeline applied for the unsupervised pipeline. Using the Pelt algorithm with a radial basis function kernel, all trajectories are ruptured into smaller, data-driven subsets (represented by the colored backgrounds), which are then clustered using a deep learning model. F) Schematic representation of the deep auto encoder used to cluster behavior in an unsupervised way. The dataset containing all instances obtained after change-point detection (*X*) is passed through a time-aware encoder consisting of a series of convolutional and recurrent layers. The output is then mapped to a bottleneck layer (*z*), in which a clustering structure is enforced by minimizing the Euclidean distance between encoder outputs and entries in a codebook, acting in practice as cluster representatives. By passing the latent embedding through the decoder, a reconstruction (*y*) is obtained for each training instance. The model is then trained to maximize the conditional log likelihood of the data, *log*(*p*(*x* | *y*)). By selecting the closest entry in the codebook for each time series segment, a sequential posterior *q*(*z*|*X*) is obtained, indicating cluster membership. As the code-lookup is non-differentiable, gradients (*∇*_*z*_ *L*) are passed through from decoder to encoder during training (red arrow in the scheme). Different colors represent different clusters. More details are available in DeepOF’s documentation.

### Time series data preprocessing

All videos and extracted time series underwent an automatic preprocessing pipeline that is included within the DeepOF module. This procedure starts by recognizing the arena from raw video by fitting an ellipse (currently, only round/elliptical arenas are supported for automatic tagging, but any polygonal arena can be defined by manual annotation using a graphical user interface (GUI); see the link in the code statement for DeepOF’s documentation details). To identify and correct any artifacts in the time series, low-quality tagging (as reported by DeepLabCut’s output likelihood) and outliers (to a fitted autoregressive model) were then marked and corrected using polynomial interpolation. In addition, smoothing was applied to the time series by cross-correlating a polynomial kernel using a Savitsky-Golay filter. Finally, coordinates were ego-centered (the cartesian origin was set to the center of each animal) and vertically aligned to remove rotational variation. A feature set was constructed as the final result of preprocessing, including the aforementioned egocentric coordinates, distances and angles between body parts, and overall locomotion speed for each mouse (Figure 1B).

### Supervised behavioral tagging with DeepOF

This feature set was used to run a supervised annotation pipeline with the objective of reporting a pre-defined set of behavioral traits on the provided videos. This set supports both dyadic interactions and individual traits, which are reported for each mouse individually (Figure 1C). Furthermore, annotated traits fall into one of two categories:

1. Traits annotated based on rules. Several motifs of interest are annotated using a few simple-to-define rules. For example, contact between animals can be reported when the distance between the involved body parts is less than a certain threshold. A comprehensive explanation of the employed rules can be found in DeepOF’s documentation (link in code statement section below). The validation of these rules using manual annotation can be found in Figure S1.
2. Traits annotated following a supervised machine learning pipeline. While rule-based annotation is enough for some traits, others are too complex or might be manifested in different subtle ways, making it difficult to detect them using this approach. For those instances, as with huddling and digging, a supervised machine learning pipeline was performed in order to offer pre-trained models within DeepOF (Figure 1D). More detailed explanations can be found in DeepOF’s documentation. The generalizability for each supervised classifier can be found in Figure S1.

Finally, a Kleinberg burst-detection algorithm (33, 34) was applied to all predictions. This step smoothens the results by merging detections that are close in time and removing isolated predictions, which an autoregressive model deems as noise.

### Unsupervised behavioral analysis with DeepOF

Unsupervised analysis of behavior was carried out using an integrated workflow within DeepOF. Starting from the afore-mentioned aligned and centered coordinates, the pipeline begins by breaking the time series of each experiment into sub-sequences in a data-driven manner. To accomplish this, DeepOF relies on the Pelt algorithm with a radial basis function kernel (35), capable of detecting significant shifts in the data in linear time (Fig 1E). Next, extracted sub-sequences for all videos were used to train a deep vector-quantization variational autoencoder model (VQ-VAE), which minimizes the mean squared error reconstruction loss between input and output, and enforces a clustering structure in the latent space using vector quantization (36). Details regarding model architecture and training can be found in DeepOF’s documentation. After training, the model is capable of clustering the input sequences by measuring the Manhattan distance of each latent embedding and the entries in a parallelly maintained codebook, selecting the latent cluster that’s most likely, given the sub-sequence at hand (Fig 1F). In practice, this unsupervised pipeline is capable of classifying the time series based on animal motion and without external information in a flexible and non-linear way. DeepOF is not the first attempt to construct an unsupervised picture of the behavioral space using autoencoders (31). Therefore, DeepOF builds on previous iterations of the idea by integrating encoding and clustering in the same model (37) using vector quantization and automating change-point detection, thus eliminating the need for the selection of a fixed behavioral time span.

### Unsupervised cluster explainability with Shapley additive explanations

In order to quantify which features might be relevant for the unsupervised models to determine the assignment of a given time segment to a given cluster, all obtained sequence-cluster mappings were analyzed using Shapley additive explanations (38, 39). First, a set of 32 distinct features describing each time chunk provided by the change-point detection algorithm in DeepOF was built. This included average speed accelerations per body part, overall locomotion speed, average triggering of all aforementioned supervised classifiers, and distance, speed, and acceleration of the spine stretch (measured as the distance between the spine 1 and tail base labels (see Figure 1A for reference). For both single animal and social interaction settings, only statistics involving the C57Bl/6N mice were utilized. Gradient boosting machines were then trained to predict cluster labels from this set of statistics after normalization across the dataset and oversampling of the minority class with the SMOTE algorithm (40). Performance was assessed by means of the area under the ROC curve across a 10-fold cross-validation loop, and feature importance was reported in terms of the average absolute SHAP values, obtained using a permutation explainer.

### Animals

8-weeks-old, in-house bred male C57Bl/6N mice were used as experimental animals. The CD1 male mice were purchased from Janvier Labs (Germany) and were used in the social avoidance and social interaction task as a social conspecific (CD1 animals were 4–6 weeks old) and as aggressors in the CSDS paradigm (CD1 animals were at least 16 weeks old). All animals were housed in individually-ventilated cages (IVC; 30cm×16cm×16cm connected by a central airflow system: Tecniplast, IVC Green Line—GM500) at least two weeks before the start of the experiment to allow acclimatization to the behavioral testing facility. All animals were kept under standard housing conditions; 12h/12h light-dark cycle (lights on at 7 a.m.), temperature 23±1°C, humidity 55%. Food (Altromin 1324, Altromin GmbH, Germany) and water were available ad libitum. All experimental procedures were approved by the committee for the Care and Use of Laboratory Animals of the government of Upper Bavaria, Germany. All experiments were in accordance with the European Communities Council Directive 2010/63/EU.

### Chronic social defeat stress

At 2 months of age, mice were randomly divided into the CSDS condition (n=27) or the non-stressed condition (n=26). The CSDS paradigm consisted of exposing the experimental C57Bl/6N mouse to an aggressive CD1 mouse for 21 consecutive days, as previously described (41). In short, the CD1 aggressor mice were trained and specifically selected on their aggression prior to the start of the experiment. The experimental mice were introduced daily to a novel CD1 resident’s territory, who attacked and forced the experimental mouse into subordination. Next, the mice were separated in the resident’s home cage using a see-through, perforated mesh and housed together overnight, allowing sensory exposure to the CD1 aggressor mouse without physical contact. Each day, for 21 consecutive days, the experimental mice were introduced to a novel CD1 aggressor mouse during a randomized moment in time between 11 a.m. and 6 p.m. to maintain unpredictability in the stress exposure and therefore avoid potential habituation. Non-stressed mice were single housed in the same room as the stressed mice. All animals were handled daily and were weighed every 3–4 days. Behavioral testing was performed after 14 days of the defeat paradigm, where behavior was observed in the morning and the defeat continued in the afternoon. The animals were sacrificed a day after the CSDS ended, which was at 3 months of age. Then, the adrenals were obtained and the relative adrenal weight was calculated by dividing the adrenal weight by the body weight before sacrifice.

### Behavioral testing

Behavioral tests were performed between 8 a.m. and 11 a.m. in the same room as the housing facility. On day 15 of the CSDS paradigm, the animals were tested on the social avoidance task, while on day 16, animals were tested on the combined open field and social interaction task. The social avoidance task was analyzed using the automated video-tracking software AnyMaze 6.33 (Stoelting, Dublin, Ireland), whereas the open field and social interaction tasks were analyzed using DeepLabCut 2.2b7 for pose estimation (27, 28), after which DeepOF module version 0.1.6 was used for preprocessing, supervised, and unsupervised analyses of behavior.

### Social avoidance

The social avoidance task was performed in the classical squared open field arena (50×50cm) to observe the social behavioral profile after CSDS, as well-established in previous studies (41, 42). The social avoidance task consisted of two phases; the non-social stimulus phase and the social stimulus phase. During the non-social stimulus phase, which was the first 2.5 minutes (min), the experimental mouse was allowed to freely explore the open field arena with a small empty wired mesh cage against the wall of the open field. Then, the empty wired mesh cage was replaced with a wired mesh cage including a trapped unfamiliar young CD1 mouse (4–6 weeks old). During the following 2.5 min, the social-stimulus phase, the experimental mouse could freely explore the arena again. The social-avoidance ratio was calculated by calculating the amount of time spent with the social stimulus, which was then divided by the time spent with the non-social stimulus.

### Open field and social interaction task

The open field and social interaction tasks were performed in a round open field arena (diameter of 38cm). The bottom of the arena was covered in sawdust material to minimize cross-over effects of stress and anxiety by the novel environment. First, the open field task was performed, during which the experimental animal was allowed to freely explore the arena for 10 min. Subsequently, for the social interaction task, an unfamiliar young CD1 (4–6 weeks old) was introduced inside the arena and both animals were allowed to freely explore the arena for 10 min. The DeepOF module can identify six behavioral traits during the single animal open field task, which include wall-climbing, digging, huddling, looking-around, sniffing, and speed (locomotion), whereas in the social interaction task, all behavioral traits can be identified (Figure 1C). During the analysis, the 10 min open field and social interaction tasks were analyzed on the total duration of the behavioral classifiers and in time bins of 2.5 min to match the time frame in the social avoidance task.

### Z-score for stress physiology and social interaction

The Z-scores combine the outcome of multiple tests via mean-normalization and provide an overall score for the related behavior of interest. Z-scores were calculated as described previously (43). The Z-score indicates for every observation (X), the number of standard deviations (σ) above or below the mean of the control group (µ). This means that for every individual observation the following formula is calculated:

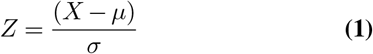

Then, the obtained values need to be corrected for the directionality, such that an increased score will reflect the increase of the related behavior of interest. This means that per test, the scores were either already correct or were adjusted in the correct directionality by multiplying with “–1”. Finally, in order to calculate the final z-score, the different z-scores per test were combined and divided by the total number of tests:

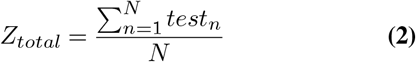

The Z-score analysis of stress physiology is based on the relative adrenal weight and the body weight at day 21 of the CSDS, which are both strongly influenced by CSDS exposure (11). The directionality of both tests did not require additional adjustment. Then, the Z-score of social interaction was calculated based on five DeepOF behavioral classifiers from the C57Bl/6N mouse, which were B-look-around, B-speed, B-huddling, B-nose-to-tail, and B-nose-to-body. The directionality was adjusted for B-speed, B-nose-to-tail, and B-nose-to-body.

### Statistics

Statistical analyses and graphs were done in RStudio (with R 4.1.1 (44)). All data were used for further statistical analysis, as no reason was observed in both the methodological as well as the statistical perspective for excluding data. Statistical assumptions were then checked, in which the data were tested for normality using the Shapiro-Wilk test and QQ-plots and for heteroscedasticity using Levene’s test. Data that violated these assumptions were analyzed using non-parametric tests. The time-course data was analyzed using the repeated measures ANOVA with time (days) as a within-subject factor and condition (non-stressed vs. stressed) as a between-subject factor. Post-hoc analysis was performed using the ANOVA test (parametric) or the Kruskal-Wallis test (non-parametric). P-values were adjusted for multiple testing using the Bonferroni method. Two-group comparisons were analyzed using independent samples t-tests (parametric), Welch’s tests (data is normalized but heteroscedastic), or Wilcoxon tests (non-parametric). Correlation analyses were performed using the Pearson correlation coefficient. The timeline and bar graphs are presented as mean ± standard error of the mean (SEM). Data was considered significant at p<0.05 (*), with p<0.01 (**), p<0.001 (***), p<0.0001 (****).

## Results

### The physiological and behavioral hallmarks of stress are reproduced by CSDS

The CSDS paradigm was performed to maintain a sustained stress exposure for several weeks, in which dysregulation of the hypothalamic-pituitary-adrenal axis (HPA-axis) and a stress-related behavioral profile were observed (Figure 2A). Animals that were subjected to CSDS showed clear hallmarks of stress exposure, as observed by a significant increase in body weight towards the end of the stress paradigm, an increase in relative adrenal weight, reduced locomotion and time spent in the inner zone of the open field, and a significantly reduced social interaction ratio in the social avoidance task (Figure 2B–F). Notably, no bodyweight difference was observed at the beginning of the CSDS paradigm (Figure 2B). In accordance with the classical analysis of the open field data (distance and inner zone time), the DeepOF module identified a reduced speed in CSDS animals (Figure S2A), and in addition, found a stress effect for look-around and sniffing (Figure S2B–C). The DeepOF behavioral classifiers wall-climbing, digging, and huddling did not show significant stress-related alterations in the open field task (Figure S2D–F).

**Fig. 2.**
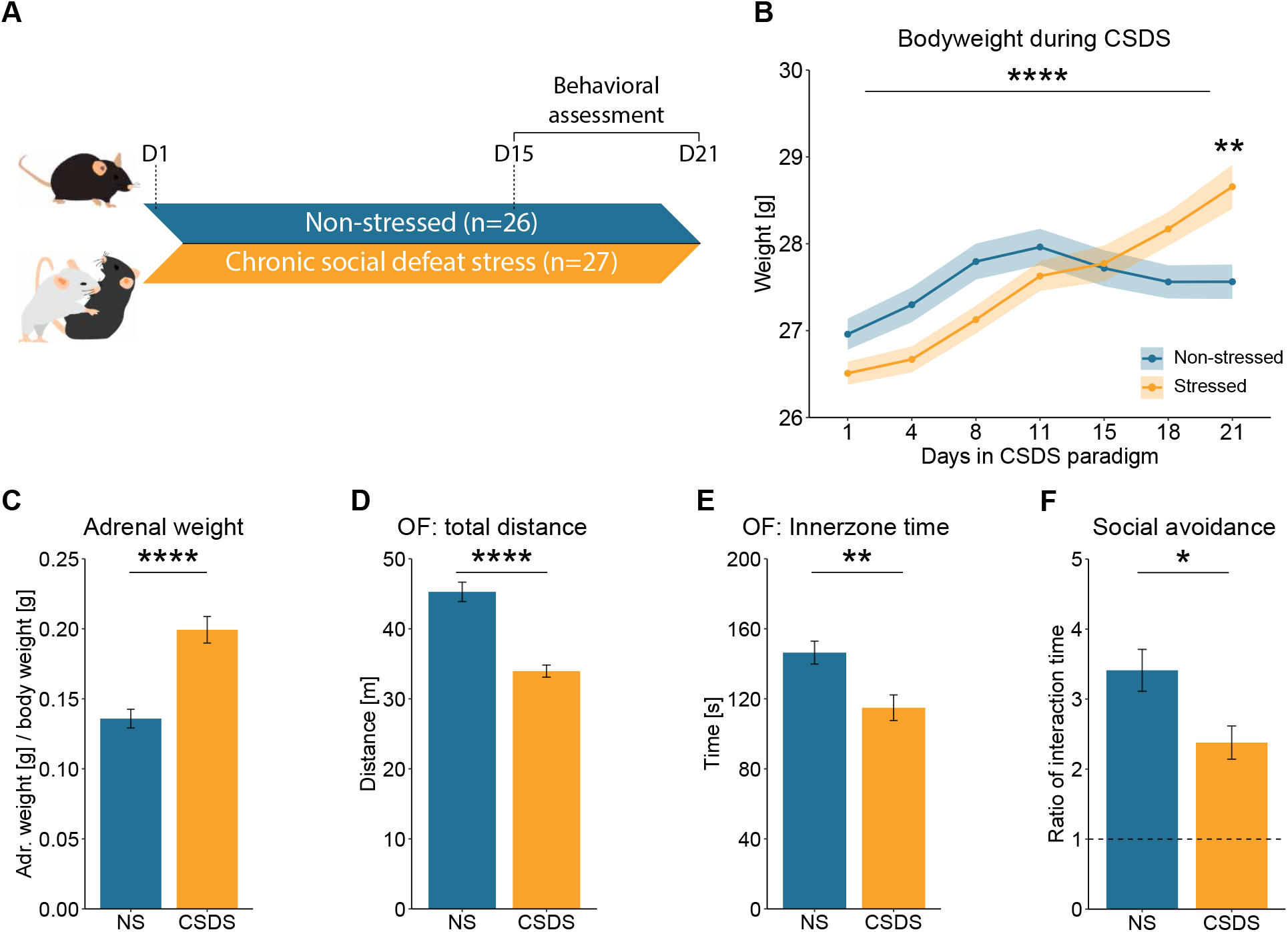
Classical hallmarks for chronic social defeat stress. A) Experimental timeline for CSDS paradigm and behavioral testing. B) Significant increase of body weight after CSDS exposure (Two-way repeated measures ANOVA: within-subject effect of time: F(6,406)=13.58, p<0.0001, as well as time×condition interaction effect: F(6,406)=6.13, p<0.0001, but no between-subject effect on condition: F(1,406)=0.20, p=0.653) & post-hoc: day 1–18 (p>0.091), day 21 (F(1,58)=11.57, p=0.007)) C) Increase of relative adrenal weight after CSDS exposure (Independent samples t-test: T(50)=–5.00, p<0.0001). D) The total locomotion in the open field (OF) was reduced after CSDS exposure (Independent samples t-test: T(51)=6.15, p<0.0001). E) The inner zone time in the open field (OF) was reduced after CSDS exposure (Independent samples t-test: T(51)=3.37, p=0.00145). F) The social interaction ratio was reduced in the social avoidance task after CSDS exposure (Wilcoxon test: W=460, p=0.0262).

### DeepOF social behavioral classifiers show a stronger PCA separation for stress exposure than social avoidance

The social behavioral pattern during the social interaction task was investigated in four non-overlapping time bins of 2.5 min each. Principal component analysis (PCA) was performed to show the difference between time bins in the social behavioral profile regardless of the animal’s stress condition (Figure 3A). Interestingly, the PCA showed a significant effect between the time bins, in which the first 2.5 min time bin was significantly different from the subsequent time bins (5, 7.5, and 10 min), whereas the subsequent time bins did not show variation between one another. (Figure 3B). This indicates that the different time bins in the social interaction task are an important variable, for which the first 2.5 min time bin needs to be especially investigated. Next, the social avoidance task and the social interaction task were compared on the strength of distinguishing between the non-stressed and stressed animals. PCAs were performed for the social avoidance task (Figure 3C) and the 2.5 min time bin social interaction data (Figure 3D–E), in which both PCAs showed a significant difference between the conditions in the principal component (PC) 1 eigenvalues (Figure 3C–E). However, the social interaction task showed a clearer separation of the conditions than the social avoidance task, indicating that the social interaction task is a more powerful tool for the identification of stressed animals compared to the social avoidance task. In addition, the PC1 top contributing behaviors for the 2.5 min time bin social interaction data were calculated using the corresponding rotated loading scores (Figure 3F). The top five contributing behaviors were reported as essential behaviors for identifying the stressed phenotype, which consisted of B-look-around, B-speed, B-huddling, B-nose-to-tail, and B-nose-to-body from the C57Bl/6N animal, whereas the other behaviors within the top 10 were either contributing to the CD1 animal (“W-” behaviors) or had a low rotated loading score (Figure 3F).

**Fig. 3.**
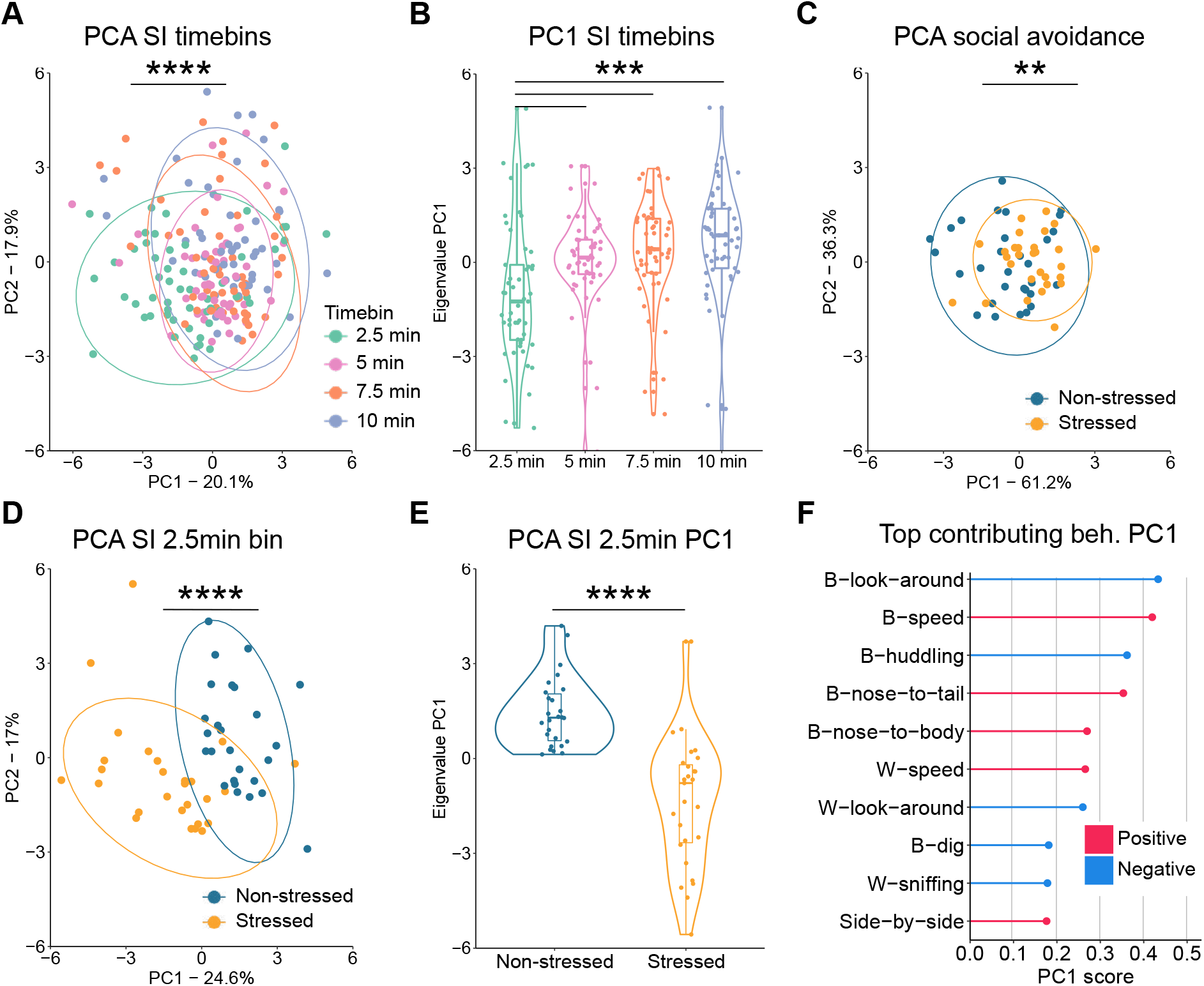
Social interaction binning yields more separable PCA projections than the social avoidance task. A) PCA revealed that the first 2.5 min time bin is significantly different from the other time bins. (One-way repeated measures ANOVA: F(3,208)=7.36, p=0.0001. B) The PC1 eigenvalues of the time bin PCA. Post-hoc: 2.5 min vs. 5 min (T(52)=–4.09, p=0.0009, 7.5 min (T(52)=–4.09, p=0.0009, 10 min (T(52)= –4.62, p=0.0002). C) The social avoidance task PCA showed a significant difference by the PC1 eigenvalues of the conditions. The PCA data consisted of the social avoidance ratio, total time spent with the non-social stimulus, and total time spent with the social stimulus. Independent samples t-test: T(57)=–2.84, p=0.006. D) The 2.5 min time bin social interaction task PCA showed a significant difference of the PC1 eigenvalues between conditions. The PCA data consisted of all the DeepOF behavioral classifiers, as listed in Figure 1C. Independent samples t-test: T(51)=6.34, p<0.0001. E) The PC1 eigenvalues of the 2.5 min time bin social interaction task. F) The top contributing behaviors of the social interaction 2.5 min time bin PC1, in which the top five behaviors were reported as the essential behaviors for identifying the stressed animals (B-look-around (–0.43), B-speed (0.42), B-huddling (–0.36), B-nose-to-tail (0.35), B-nose-to-body (0.27). “B-” indicates C57Bl/6N behaviors and “W-” indicates CD1 behaviors.

### DeepOF social behavioral classifiers are strongly altered by CSDS

Next, the influence of the CSDS on the top five contributing behaviors in the social interaction task was investigated. At first, the total duration of the social interaction task was analyzed, regardless of the time bins. Although, the behavioral effects for B-look-around and B-speed from the C57Bl/6N animal on CSDS exposure were already apparent, in the majority of the top contributing behaviors, no significant CSDS effect could be observed when analyzing the full 10 min (Figure S3, barplots). However, when looking at the total duration per time bin for the top contributing behaviors, the 2.5 min time bin data show a clear CSDS effect (Figure S3, timelines), whereas the 5-, 7.5-, and 10-min time bins do not show a CSDS effect, in accordance with the PCA time bin analysis. More specifically, the 2.5 min time bin analysis showed that the CSDS exposure significantly elevated the duration for B-look-around (Figure 4A) and B-huddling (Figure 4B), while the B-speed (Figure 4C), B-nose-to-tail (Figure 4D), and B-nose-to-body (Figure 4E) were significantly lowered. In addition, Figure S4 shows the total duration for all other DeepOF behavioral classifiers, in which a significant stress effect is observed for B-wall-climbing (Figure S4E).

**Fig. 4.**
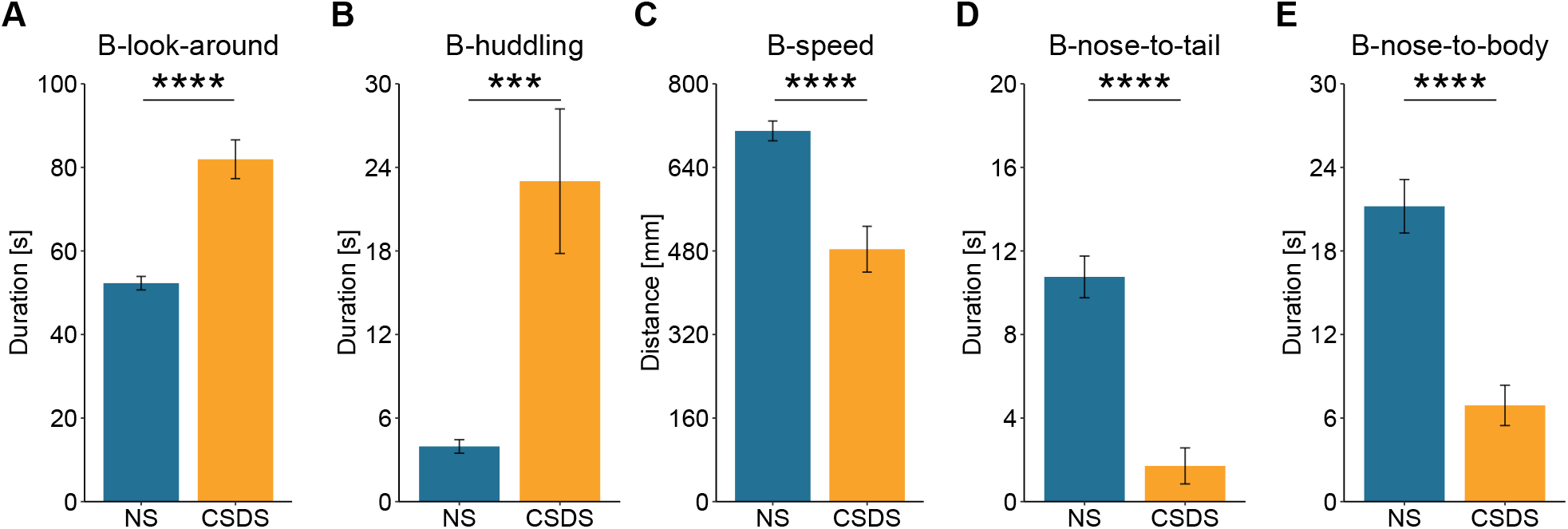
Top contributing behaviors of the 2.5 min time bin social interaction task. A–B) B-look-around and B-huddling duration were significantly elevated in stressed animals. Wilcoxon tests: W_A_=56, p_A_<0.0001 and W_B_=145.5, p_B_=0.0003, respectively. C–E) B-speed, B-nose-to-tail, and B-nose-to-body duration were significantly lowered in stressed animals. Wilcoxon tests: W_C_=573, p_C_<0.0001, W_D_=664, p_D_<0.0001, and W_E_=635, p_E_<0.0001, respectively.

### Z-score for DeepOF social interaction correlates with Z-score for stress physiology

The Z-score of stress physiology was calculated using the relative adrenal weight and body weight on day 21 of the CSDS. The stress physiology Z-score provides a strong CSDS profiling tool and was used for correlation analysis between the social avoidance and social interaction tasks. No significant correlation was observed between the Z-score of stress physiology and the social avoidance ratio (Figure 5A). Subsequently, the Z-score of social interaction was calculated by using the 2.5 min time bin of the top five contributing behaviors in the social interaction task (Figure 4A–E). Stress physiology and social interaction Z-score showed a significant positive correlation (Figure 5B), which indicates that the social interaction Z-score provides a stronger tool for CSDS profiling compared to the social avoidance ratio. Next, correlation analyses were performed between the Z-score of social interaction and all other behavioral and physiological measurements which indicated, a strong correlation with several open field parameters, such as distance and inner zone entries, but interestingly no correlation with the social avoidance ratio (Figure 5C).

**Fig. 5.**
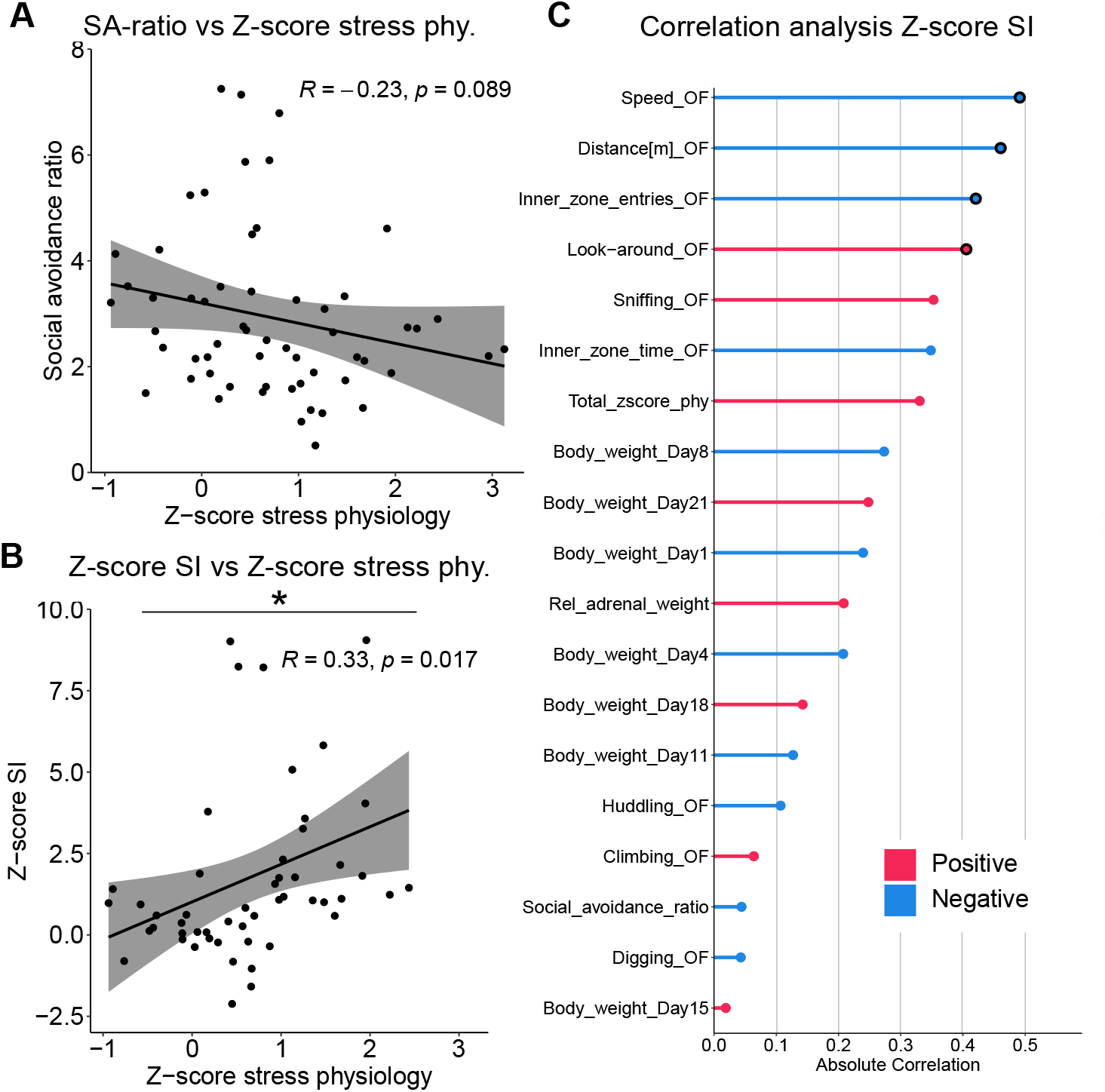
Z-score correlation analysis for social interaction task and social avoidance task. A) Pearson correlation analysis between the social avoidance ratio and the Z-score of stress physiology. Pearson correlation: R=–0.23, p=0.089. B) Pearson correlation analysis between the social interaction task 2.5 min time bin top five contributing behaviors and the Z-score of stress physiology. Pearson correlation: R=0.33, p=0.017. C) Pearson correlation analyses between the Z-score of social interaction and all other parameters. A strong correlation was observed with several open field parameters, such as distance (R=–0.46, p=0.0005) inner zone: entries (R=–0.42, p=0.0017), but no correlation was found with the social avoidance ratio (R=–0.044, p=0.76). Black circles around the points are identified as significant correlations, p<0.05.

### DeepOF recognizes overall differences in behavior in an unsupervised way

When applying the VQVAE-based unsupervised pipeline within DeepOF, the number of clusters is a hyperparameter the user must tune. However, an optimal solution can be found by selecting the number of clusters that explain the largest difference between experimental conditions (in terms of the area under the ROC curve of a linear classifier to distinguish between them). This procedure yielded an optimal of 12 clusters for the social interaction task and 9 for the single animal setting (Figure 6A and Figure S5A). Once the number of clusters was fixed, the stress-induced phenotype was investigated across time. Therefore, a growing time window spanning an increasing number of sequential seconds was analyzed. For each analysis, the discriminability between conditions was tested by evaluating the performance of a linear classifier to distinguish between them in an aggregated embedding space, for which each experiment is represented by a vector containing the time spent per cluster (Figure 6B and Figure S5B, grey curves). The bin size for which discriminability was maximized was then selected as optimal and continuously used for further analysis. In this case, we observed an optimum of 2.06 minutes for the social interaction task, indicating that differences between conditions are maximized early in the 10-minute-long experiments. Furthermore, performance across consecutive, non-overlapping bins retaining the optimal size was also reported (Figure 6B and Figure S5B, dark green curves). Here, decaying performance across bins in the social interaction setting is compatible with a state of arousal, where conditions become less distinguishable over time after the behavior of the C57Bl/6N mice becomes less influenced by novelty. The largest difference between stressed and non-stressed animals can thus be observed during this period. In line with this finding, the optimal distance in the single animal data was reached at 8.68 minutes, suggesting that no binning is necessary since behavior between conditions remains consistently distinguishable across the videos (Figure S5B).

**Fig. 6.**
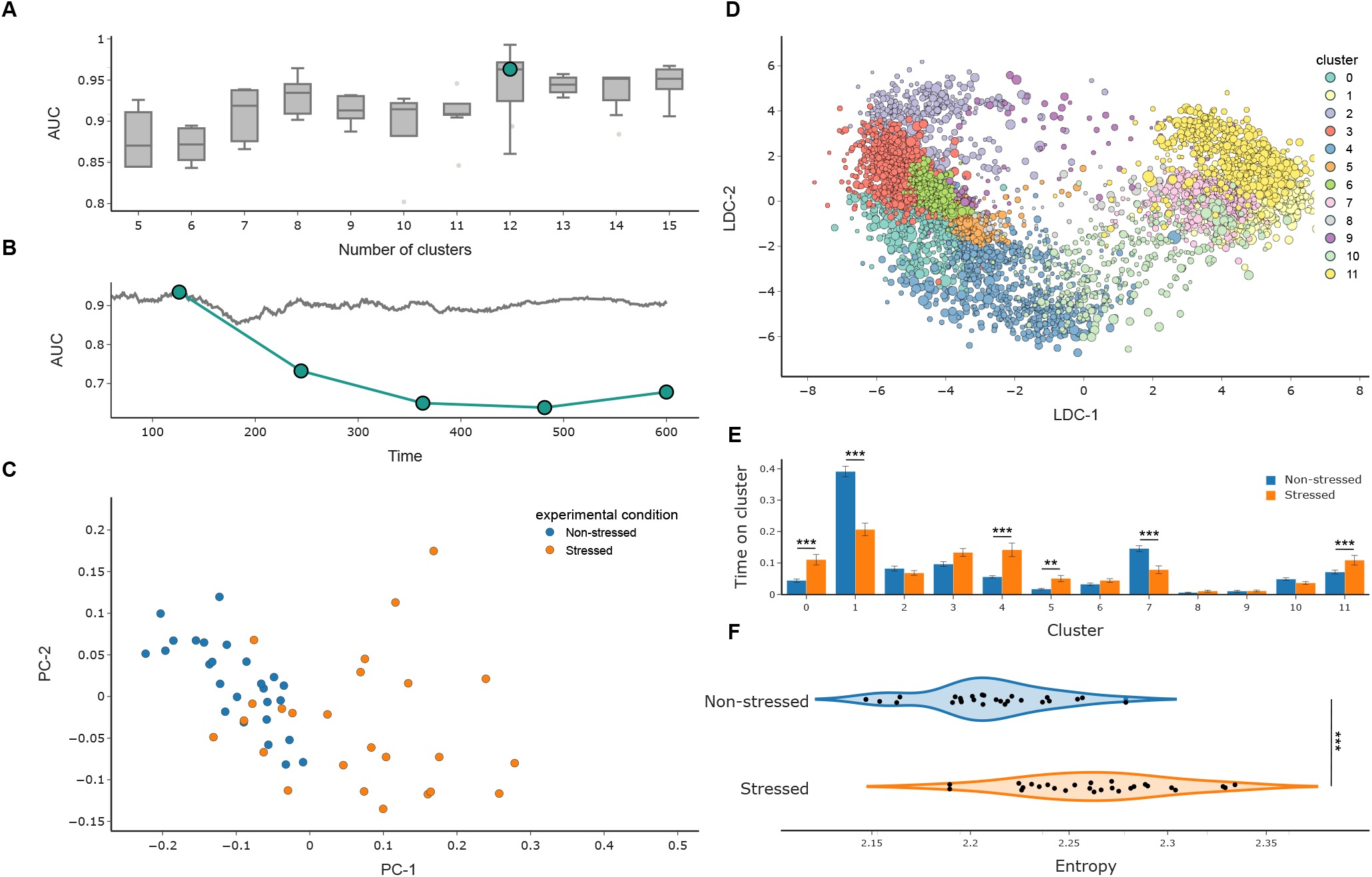
VQVAE unsupervised analyses identify different behavioral patterns between stressed and non-stressed mice during the social interaction task. A) Cluster selection pipeline results. Models ranging from 5 to 25 clusters were trained in a 10-fold cross-validation loop (only 5 to 15 are shown for visualization purposes). Discriminability between conditions on the aggregated embeddings (analogous to panel B) is reported. The cluster number which maximized this curve (explaining the largest difference between the conditions of interest) was selected. B) Optimal binning of the videos was obtained by measuring the performance of a linear classifier on discriminating between both conditions across a growing window, taking a range from the first 60 to 600 seconds for each video (grey curve). Higher values correspond to more distinguishable distributions (that is larger differences in behavioral profiles across conditions). A maximum was observed at 2.06 minutes, close to the stipulated 2.5 minutes selected based on the social avoidance task. The dark green curve depicts discriminability across all subsequent bins of optimal length (first 2.06 minutes, second 2.06 minutes, etc.). The decay observed across time corresponds to the hypothesized arousal period in the CSDS cohort. C) Representation of the aggregated unsupervised embedding for the optimally discriminant bin per experimental video colored by condition. Each 10 min video was fed to a trained VQVAE model using DeepOF’s unsupervised pipeline. Aggregated embeddings were constructed as the time spent on each defined cluster (yielding a 12-dimensional vector). Dimensionality was further reduced using PCA for visualization purposes. D) Unsupervised embedding of all automatically ruptured video fragments. Different colors correspond to different clusters as recognized by DeepOF. Dimensionality was further reduced using LDA for visualization purposes. E) Cluster population per experimental condition in the first optimal bin (2.06 min). Reported statistics correspond to a 2-way Welch t-test corrected for multiple testing using Bonferroni’s method across both clusters and bins. F) Stationary distribution’s entropy of the transition matrices provided by the unsupervised pipeline within DeepOF. Higher values suggest that stressed animals explore the behavioral space in a more erratic way (t-test p-value=6.6 ×10^−7^).

Moreover, the aforementioned aggregated embedding per experimental C57Bl/6N animal was capable of representing the difference between non-stressed and stressed animals in both social interaction and single animal settings (Figure 6C and Figure S5C). However, the stress-induced difference was substantially greater during the social interaction task than during the open field task (areas under the ROC curve for a linear classifier to distinguish between conditions of 0.921 and 0.837, respectively).

### Individual unsupervised clusters reveal differences in both behavior enrichment and dynamics across conditions

Going beyond global differences in behavior, the aggregated embeddings were the result of summarizing the expression of a set of detected behavioral clusters (Figure 6D and Figure S5D). Once obtained, DeepOF allows comparing both their enrichment (or differential expression) and dynamics between conditions. In terms of enrichment, the time on each cluster across all videos for each condition is recorded in the optimal bin. Importantly, DeepOF has no knowledge of which video the processed sequence belongs to, and therefore does not know its condition at the time of training and assigning clusters. Expression was then compared using 2-way Kruskal-Wallis tests for each cluster independently, and p-values were corrected for multiple testing using Bonferroni’s method. We observed significant differences in 7 out of 12 and 6 out of 9 clusters for social interaction and single animal data, respectively (Figure 6E and Figure S5E). In terms of dynamics, an empirical transition matrix was obtained for each condition by counting how many times an animal goes from one given cluster to another (including itself). Since all transitions were observed to have non-zero probability, the Markov chains obtained from simulations can be proven to reach a steady-state over time (where probabilities to go from one behavior to another stabilize). The entropy of these steady-state distributions was reported for both conditions, with higher values corresponding to a less predictable exploration of the behavioral space. Interestingly, stressed animals showed a significantly higher steady-state entropy in the social interaction task than their non-stressed counterparts (Kruskal-Wallis p-value = 6.64×10-7), suggesting more erratic behavior, which could be explained by an increased susceptibility which is influenced by the CD1 counterparts (Figure 6E). In line with this hypothesis, the single animal experiments show no significant difference (Figure S5E).

### Shapley additive explanations reveal a consistent profile across differentially expressed clusters

An important aspect of machine learning that is relevant for highly complex models is its explainability. In this study, we aimed to explain cluster assignments by fitting supervised classifiers (gradient boosting machines) that map statistics of the initial time series segments (including locomotion and individual body part speeds, distances, and angles) to the subsequent cluster assignments. Performance and generalizability of the constructed classifiers across the dataset were assessed in terms of the area under the ROC curve on a 10-fold stratified cross-validation loop, which was designed so that segments coming from the same video were never assigned to both train and test folds. Data were standardized, and the minority class was oversampled using the SMOTE algorithm to correct for class imbalance. The result of this analysis is a set of feature explainers for each retrieved cluster, which can be used to interpret, alongside visual inspection of the video fragments in a cluster, what the obtained behavioral motifs represent. In this context, we found consistent descriptions of clusters enriched in stressed animals for the social interaction task (Figure 7A-E). For example, social interaction clusters 0, 4, and 5 are negatively associated with locomotion speed and spine stretch, corresponding to more passive behaviors. Upon visual inspection, we identified cluster 0 as passive huddling not necessarily accompanied by social interaction and cluster 5 as enriched in segments where the C57Bl/6N animals are being actively approached by their counterparts. In contrast, clusters 1 and 7 (enriched in non-stressed animals) represent more active, locomotion-engaged roles. These patterns can be observed as well in the single-animal data (Figure S6A–D).

**Fig. 7.**
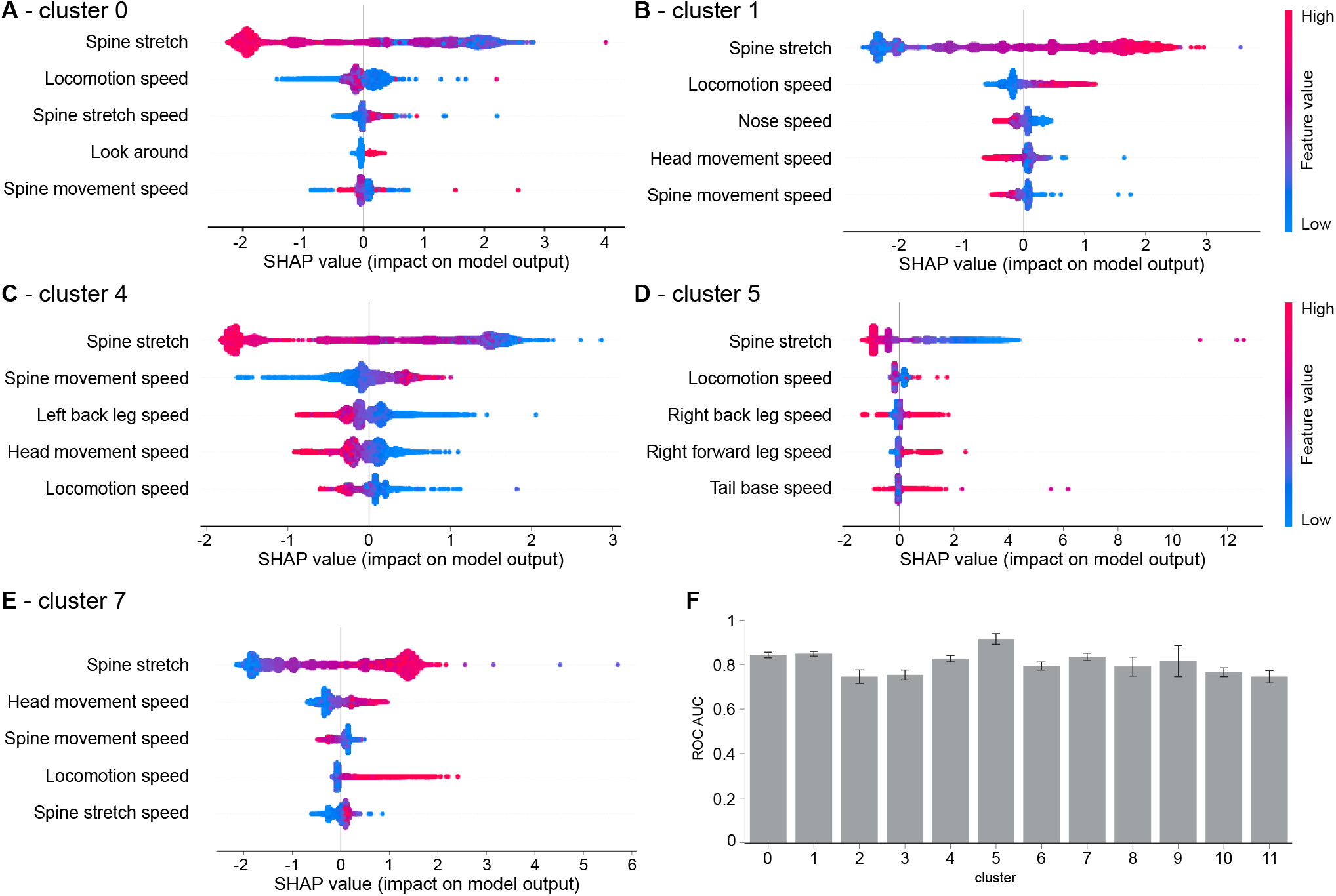
SHAP analysis of unsupervised cluster assignments in the social interaction task. A–E) Beeswarm plots for clusters 0, 1, 4, 5, and 7 identified with the unsupervised DeepOF pipeline on the social interaction experiments. Gradient boosting machines were trained to map from a predefined set of time series statistics (including locomotion and individual bodypart speeds, distances, and angles) to the previously obtained cluster assignments. The depicted plots display the first 5 most important features for each classifier, in terms of the mean absolute value of the Shapley additive values (SHAP). F) Performance of the constructed gradient boosting classifiers in terms of the test area under the ROC curve. Data were standardized and the minority class was oversampled using the SMOTE algorithm to correct for class imbalance.

The performance of all gradient boosting machines when predicting cluster membership in a 10-fold cross-validation fashion can be found in Figure 7F (for the social interaction experiments) and Figure S6E (for the single-animal experiments). In addition, representative clips of all clusters for both social interaction and single animal data can be found in the supplemental material (see data availability statement below).

## Discussion

For decades there has been a trend to standardize and simplify social behavioral tests, which has led to an oversimplification of the social behavioral repertoire. The current developments of open-source markerless pose estimation tools for tracking multiple animals have provided the possibility for more complex and socially relevant behavioral tests. The current study provides an open-source tool, “DeepOF”, which can investigate both the individual and social behavioral profile in mice using DeepLabCut-annotated pose estimation data. The current study identified a distinct social behavioral profile following CSDS using a selection of five DeepOF social behavioral classifiers from the C57Bl/6N animal, consisting of B-look-around, B-speed, B-huddling, B-nose-to-tail, and B-nose-to-body. In addition, a similar social behavioral profile was identified with the unsupervised workflow, which was capable of detecting behavior enrichment and dynamic differences across conditions — especially during the social interaction task, but also in single animal open field data. Next, we identified the first minutes during the interaction with a novel conspecific as crucial for the social profiling of CSDS exposure in both supervised and unsupervised workflows. Overall, this study demonstrates the high utility and versatility of DeepOF for the analysis of complex individual and social behavior in rodents.

### DeepOF as part of a markerless pose estimation toolset

The initial release of DeepLabCut in 2018 (27) provided a reliable and accessible tool for researchers around the globe to process markerless pose estimation data, which has undoubtedly changed the field of behavioral neuroscience. This has set in motion a rapid growth of tools for analyzing pose estimation data that are increasing the range of possibilities in the field, which were unimaginable using classical tracking approaches or manual scoring. An important distinction between these pose estimation analysis tools is whether they intend to extract pre-defined and characterized traits (supervised) or to explore the data and extract patterns without external information (unsupervised). The DeepOF module is designed to provide both analysis pipelines; the supervised behavioral classifiers offer a quick and easy-to-use analysis to detect individual and social behavioral traits. In addition, when differences between the conditions are not reflected in these traits, or the researcher aims to obtain behavioral embeddings, the DeepOF module can encode the data in a time-aware way that can report differentially expressed patterns in an unsupervised manner.

### The supervised framework: spotting recognizable patterns

The supervised pipeline within the DeepOF package can be used on single open field and dyadic social interaction behavioral data. DeepOF is capable of reporting a pre-defined set of behavioral traits without any extra labeling or training required. In order to accomplish this, it relies on simple rule-based annotations and machine learning binary classifiers whose generalizability has been tested, trading off flexibility for ease of use. This makes it user-friendly for researchers without computational expertise to apply this supervised pipeline, without having to make any modifications. To further detect, unsupported patterns, using a more involved and flexible tool (such as SimBA (34) or MARS (26)) could be a reasonable next step to take. These tools include a supervised approach that requires the user to label and train classifiers, providing the freedom to train powerful classifiers and recognize behavioral traits, which is especially beneficial for labs without computational expertise. However, in contrast to DeepOF, this approach also delegates to the user the responsibility of testing the generalizability of the results (how well the trained models can be applied to newly generated data, even in similar settings), which requires careful practices from the experimenters.

### The DeepOF module provides a more complete social behavioral profile than the social avoidance task

The social behavioral profile in CSDS-subjected animals has been measured extensively using the social avoidance task, which is based on the separation of social behavioral traits between stressed and non-stressed animals (10, 15, 45). Previous research has shown that rodents have a social interaction preference towards a novel conspecific compared to a familiar conspecific (46). However, the duration of this social behavioral arousal state has not been well documented. In this context, and by replicating the time the social avoidance task typically lasts for (9), the current study shows that the CSDS-related social behavioral profile, obtained with the DeepOF supervised classifiers, was increasingly observed during the first 2.5 min of the 10 min social interaction task. Furthermore, the presented unsupervised workflow was used to determine an optimal binning of our experiments by measuring how different both conditions were across time for a linear classifier. This yielded an optimal separation at approximately two minutes, which then decayed over subsequent time bins in a manner consistent with the arousal hypothesis. The fact that this result was not seen in the absence of a conspecific strengthens this argument. Taking this into account, we argue that the introduction of a novel conspecific induces a state of arousal, which coincides with a distinct social behavioral profile that disappears over time after 2-to-3 minutes due to habituation. Along these lines, this study shows that the DeepOF social behavioral classifiers provide a stronger separation of the social behavioral profile between stressed and non-stressed animals compared to the classical social avoidance task. In addition, the Z-score of DeepOF social behavioral classifiers showed a significant correlation with the Z-score of stress physiology, while the social avoidance data did not correlate with the Z-score of stress physiology. Taken together, it can be concluded that using the DeepOF social behavioral classifiers provides a more robust and clearer social behavioral profile in animals subjected to CSDS compared to the social avoidance task. An important reason for the superiority of DeepOF in social behavioral profiling depends on the experimental set-up: the social avoidance task relies on the confinement of an animal (for example using a wired mesh cage), which means that no natural interaction between freely-moving animals is possible, whereas the social interaction task is based on a naturally occurring interaction between freely moving animals (16). Moreover, in the social avoidance task, the confined animal can show symptoms of anxiety-related behavior, which influences the physiological state and the social interaction and approach behavior of the conspecific (47–49). An important advantage of the DeepOF module is the many different behavioral classifiers that can be investigated at the same time without increasing the labor intensity. The combined analysis of multiple behavioral classifiers into a Z-score of social behavior provides a more complete social behavioral profile than solely investigating social avoidance behavior, which may also be beneficial for improved classification of identifying stress susceptible and resilient individuals.

### DeepOF can detect and explain differences across conditions even when no labels are available

The supervised pipeline within DeepOF follows a highly opinionated philosophy, which focuses on ease of use and relies on predefined models. As an alternative, DeepOF offers an unsupervised workflow capable of representing animal behavior across experiments without any label information. In its most basic expression, this involves obtaining a representation for each experiment in a time-aware manner: unlike other dimensionality reduction algorithms like PCA, UMAP, and T-SNE (22), DeepOF, when applied to the raw dataset, relies on a combination of convolutional and recurrent neural networks capable of modeling the sequential nature of motion. Each input to the models consists of a subsequence across a non-overlapping sliding window of each experiment. Although this idea has been explored before (31), DeepOF has the advantage of detecting automatically where each experimental sequence needs to be split, which is based on a change point detection algorithm that detects when the animals change their behavior. In addition, these global embeddings can be decomposed into a set of clusters representing behavioral motifs that the user can then inspect both visually and with machine learning explainability methods. Moreover, by comparing cluster enrichment and dynamics across conditions, it is possible to answer questions that are relevant towards understanding what the observed difference might be based on, without any previous knowledge: which behaviors are most or least expressed in each condition? Is the set of behaviors explored differently in non-stressed and stressed mice? This constitutes a complementary approach that can be beneficial to further direct hypotheses when little knowledge is available. In addition, by not only showing overall differences between cohorts but also reporting which motion primitives might be driving them, it is possible to test hypotheses by training novel supervised classifiers based on those motion primitives. This will allow researchers to distinguish new, meaningful patterns that have not been reported before and that may be significantly associated with a given condition. Taken together, the current study shows that the DeepOF unsupervised pipeline does not only recapitulate results previously obtained with the supervised analysis, but also shows how this tool can be used to detect habituation and overall differences in behavioral exploration. We also show that detected differences are significantly stronger when a conspecific is present, although also detectable during single animal open field exploration.

### Towards an open-source behavioral analysis ecosystem

One of the main advantages of DeepOF, SimBA (19), VAME (31), MARS (26), and many other packages cited in this manuscript, is that they are open source. This means that how these tools are functioning is transparent and it is possible for the community to contribute to their development. We strongly believe that the adoption of open-source frameworks can not only increase transparency in the field but also incentivize a feeling of community, in which researchers and developers can share ideas, code, and solve bugs and problems together. Moreover, open-source facilitates beneficial feed-back loops, where the data generated using these tools can be published, thus increasing the opportunity to produce better software. A good example of this being zero-shot pose estimation (50), which enables motion tracking without labeling, by cleverly leveraging information from several publicly available datasets. In addition, new technologies are starting to enable joint learning from multiple modalities, such as neural activation and behavior (51), which opens the door to model how one affects the other. Finally, in contrast to several other options that offer extended functionality but rely on proprietary algorithms and/or specialized hardware (23), these tools have the potential to make otherwise expensive software available to a larger audience.

## Conclusion

Taken together, the current study provides a novel approach for individual and social behavioral profiling in rodents by both extracting pre-defined behavioral classifiers and using unsupervised, time-aware embeddings using DeepOF. We show evidence for the validation of the behavioral classifiers and provide an open-source package in order to increase transparency and contribute to the further standardization of the behavioral constructs. We also show that, while differences across conditions are detectable during single animal open field exploration, they are enhanced in the social interaction task involving a companion mouse. Furthermore, while the classical social avoidance task does identify the social behavioral profile induced by CSDS, the DeepOF behavioral classifiers provide a more robust and clearer profile. In addition, the first minutes of the interaction with a novel conspecific coincide with a state of social arousal, which disappears with time due to habituation. This study shows that the DeepOF module can contribute to further unraveling the role of social and non-social behaviors in stress-related disorders, such as MDD. In addition, the DeepOF module contributes to a more specific classification of the affected individual- and social behaviors in stress-related disorders, which could contribute to the study of drug development for psychiatric disorders.

## Author contributions

JB and MVS conceived the study. LM wrote the DeepOF package, with primary technical assessment from FA, BP, and BMM. JB and MR performed the experiments. LMB, LvD, and CE assisted with the experiments. JB and LM analyzed the data and wrote the first version of the manuscript. BP worked on figure design. All authors contributed to the revision of the manuscript.

## Code availability statement

All data and the accompanying code to perform the analyses and creating the figures are available via a [GitLab](https://gitlab.mpcdf.mpg.de/lucasmir/deepof) repository. Documentation is available on [read the docs](https://deepof.readthedocs.io/en/latest/).

## Funding

This study is supported by the “Kids2Health” grant of the Federal Ministry of Education and Research [01GL1743C], and the European Union’s Horizon 2020 research and innovation programme under the Marie Skłodowska-Curie grant agreement No. 813533.

## ACKNOWLEDGEMENTS

The authors thank the DeepLabCut development team for creating and maintaining the DeepLabCut software. In addition, the authors thank Sowmya Narayan for the language editing of the manuscript.

## Supplemental material

**Fig. S1.**
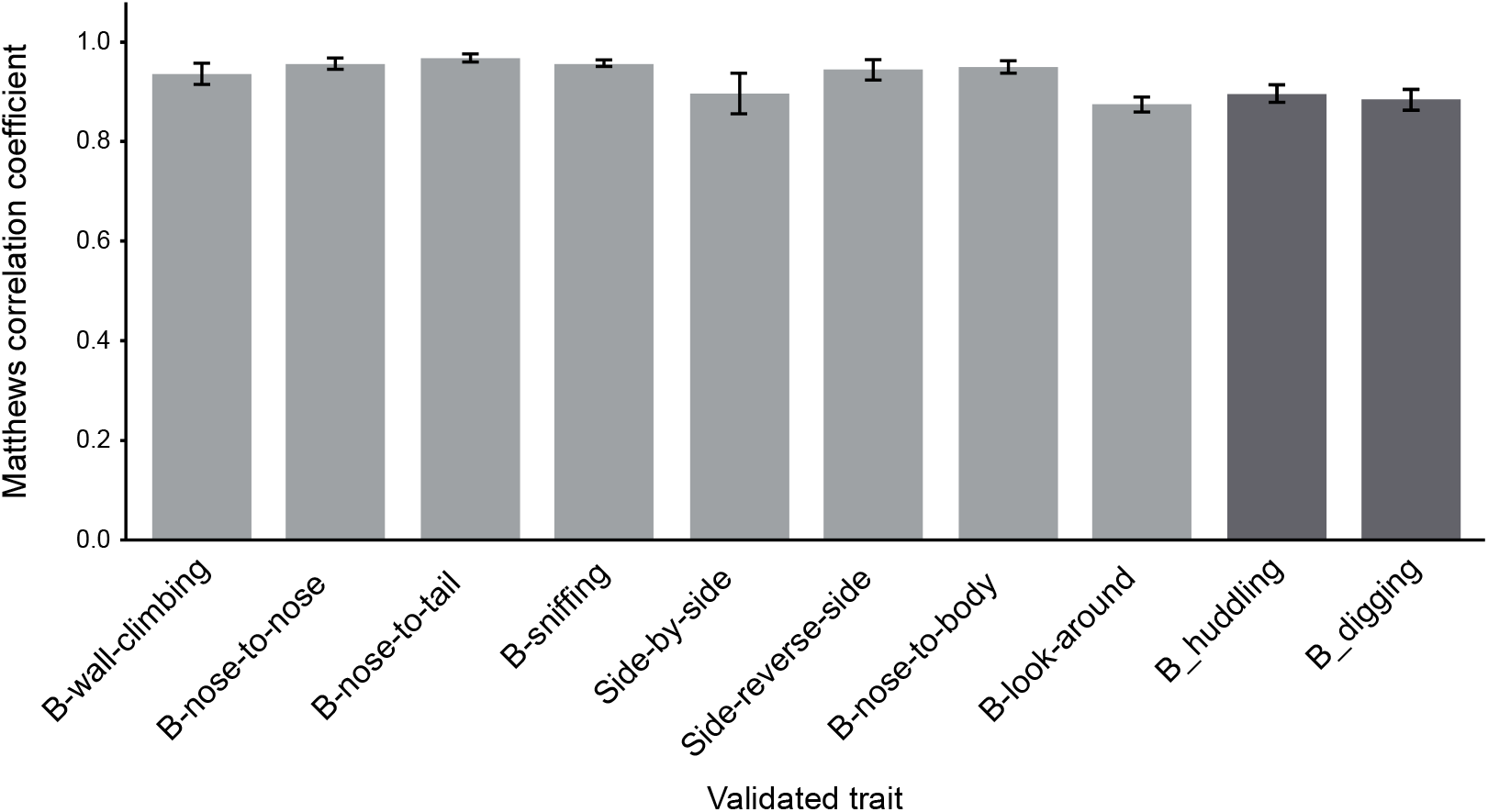
Validation of annotated behaviors. 10 out of 53 videos were manually annotated for all traits using the Colabeler software v2.0.4. The Matthews correlation coefficient between manual labels and predicted binary outcomes (presence or absence of a given trait at a given time) is reported, where 0 and 1 correspond to random and perfect predictions, respectively. Error bars represent the standard error of the mean across experiments. Dark blue bars represent traits that were annotated using simple rules, whereas orange bars represent traits that were annotated using supervised machine learning models (trained using a 5-fold nested cross-validation approach – hyperparameters tuned using 5×5-fold nested cross-validation).

**Fig. S2.**
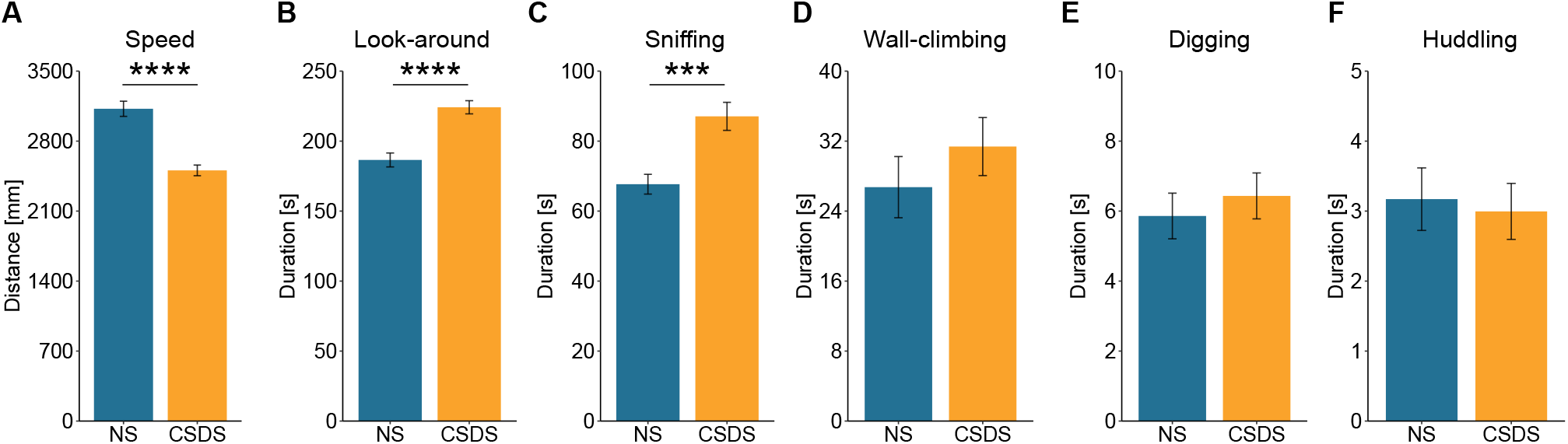
DeepOF behavioral classifiers in the open field task. A) Speed is significantly lower in stressed animals. Independent samples t-test: T(51)=6.69, p<0.0001. B) Look-around is significantly higher in stressed animals. Independent samples t-test: T(51)=–5.50, p<0.0001. C) Sniffing is significantly higher in stressed animals. Independent samples t-test: T(51)=–3.93, p=0.00026. D) Wall-climbing is not significantly different between stressed and non-stressed animals. Wilcoxon test: W=283.5, p=0.233 E) Digging is not significantly different between stressed and non-stressed animals. Independent samples t-test: T(51)=–0.62, p=0.538. F) Huddling is not significantly different between stressed and non-stressed animals. Wilcoxon test: W=369.5, p=0.749.

**Fig. S3.**
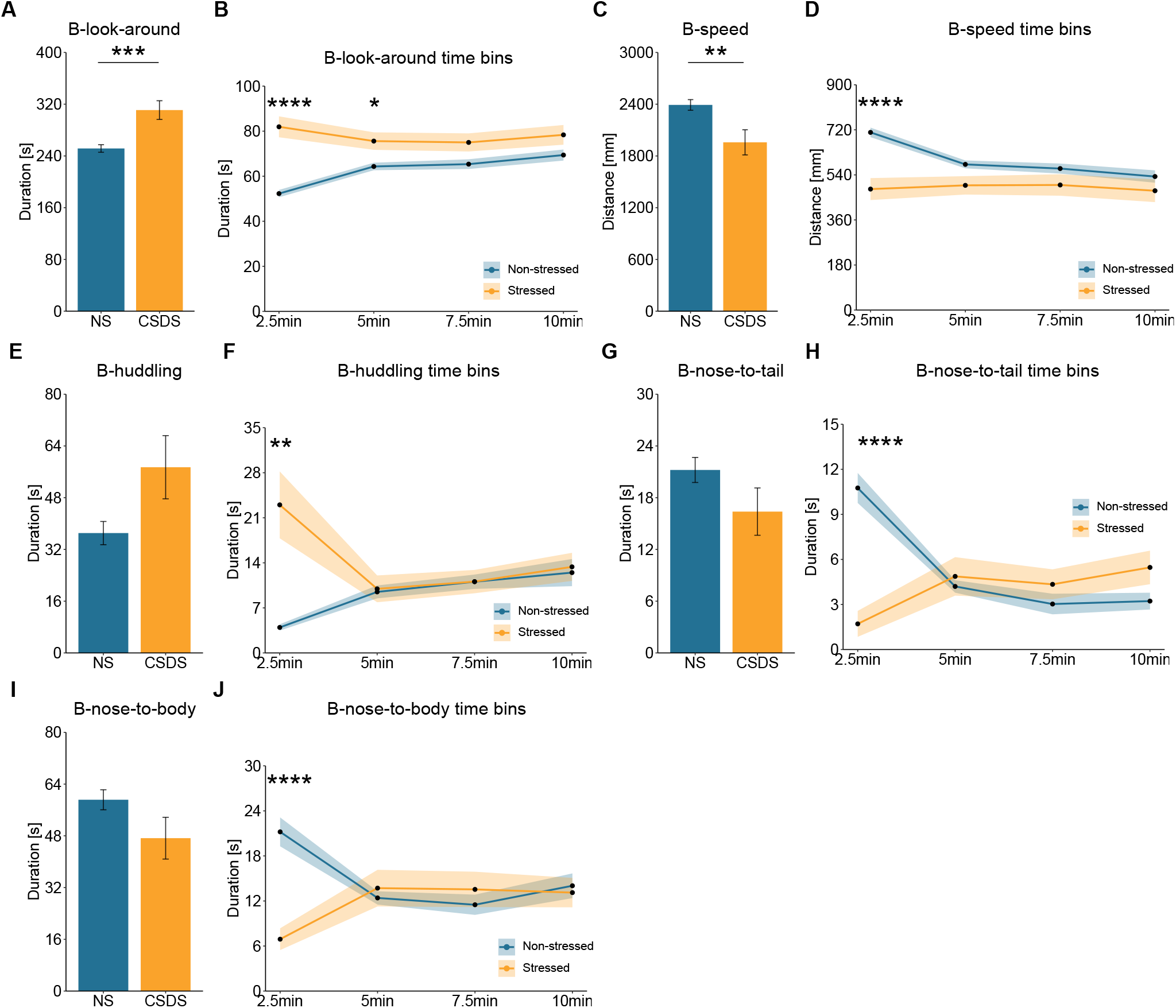
Top five contributing behaviors in the social interaction task for 10 min total duration and time bins. A) The total duration B-look-around is significantly higher in stressed animals. Welch’s test: T(34.22)=–3.81, p=0.0006. B) B-look-around is significantly higher in stressed animals for the 2.5 min time bin (T(51)=35.33, p<0.0001) and the 5 min time bin (T(51)=6.95, p=0.044), but not for the 7.5 (T(51)=4.39, p=0.164) and 10 (T(51)=3.14, p=0.328) min time bins. (Two-way repeated measures ANOVA: condition effect: F(1,208)=39.33, p<0.0001, no time effect: F(1,208)=3.52, p=0.062, but an interaction effect condition×time: F(1,208)=9.01, p=0.003). C) The total duration B-speed is significantly lower in stressed animals. Welch’s test: T(34.95)=2.74, p=0.0095. D) The time B-speed is significantly lowered in stressed animals for the 2.5 min time bin (T(51)=21.89, p<0.0001), but not for the 5 (T(51)=4.29, p=0.172), 7.5 (T(51)=1.88, p=0.708) and 10 (T(51)=1.19, p=1) min time bins. (Two-way repeated measures ANOVA: condition effect: F(1,208)=20.92, p<0.0001, time effect: F(1,208)=6.87, p=0.009 and an interaction effect condition×time: F(1,208)=6.21, p=0.013). E) The total duration B-huddling does not show a significant difference between stressed and non-stressed animals. Wilcoxon test: W=302, p=0.388. F) The time B-huddling is significantly higher in stressed animals for the 2.5 min time bin (H(1)=13.37, p=0.0010), but not for the 5 (H(1)=1.49, p=0.892), 7.5 (H(1)=0.86, p=1) and 10 (H(1)=0.02, p=1) min time bins. (Two-way repeated measures ANOVA: condition effect: F(1,208)=8.46, p=0.004, no time effect: F(1,208)=0.011, p=0.916, but an interaction effect condition×time: F(1,208)=12.22, p=0.0006). G) The total duration B-nose-to-tail does not show a significant difference between stressed and non-stressed animals. Welch’s test: T(39.38)=1.56, p=0.128. H) The time B-nose-to-tail is significantly lowered in stressed animals for the 2.5 min time bin (H(1)=31.59, p<0.0001), but not for the 5 (H(1)=3.14, p=0.31), 7.5 (H(1)=0.032, p=1), and 10 (H(1)=0.59, p=1) min time bins. (Two-way repeated measures ANOVA: condition effect: F(1,208)=3.27, p=0.072, time effect: F(1,208)=4.31, p=0.039 and an interaction effect condition×time: F(1,208)=33.52, p<0.0001). I) The total duration B-nose-to-body does not show a significant difference between stressed and non-stressed animals. Welch’s test: T(37.26)=1.66, p=0.105. J) The time B-nose-to-tail is significantly lowered in stressed animals for the 2.5 min time bin (H(1)=25.54, p<0.0001), but not for the 5 (H(1)=0.86, p=1), 7.5 (H(1)=0.026, p=1) and 10 (H(1)=0.45, p=1) min time bins. (Two-way repeated measures ANOVA: condition effect: F(1,208)=5.01 p=0.024, no time effect: F(1,208)=0.075, p=0.785, but an interaction effect condition×time: F(1,208)=11.92, p=0.0007).

**Fig. S4.**
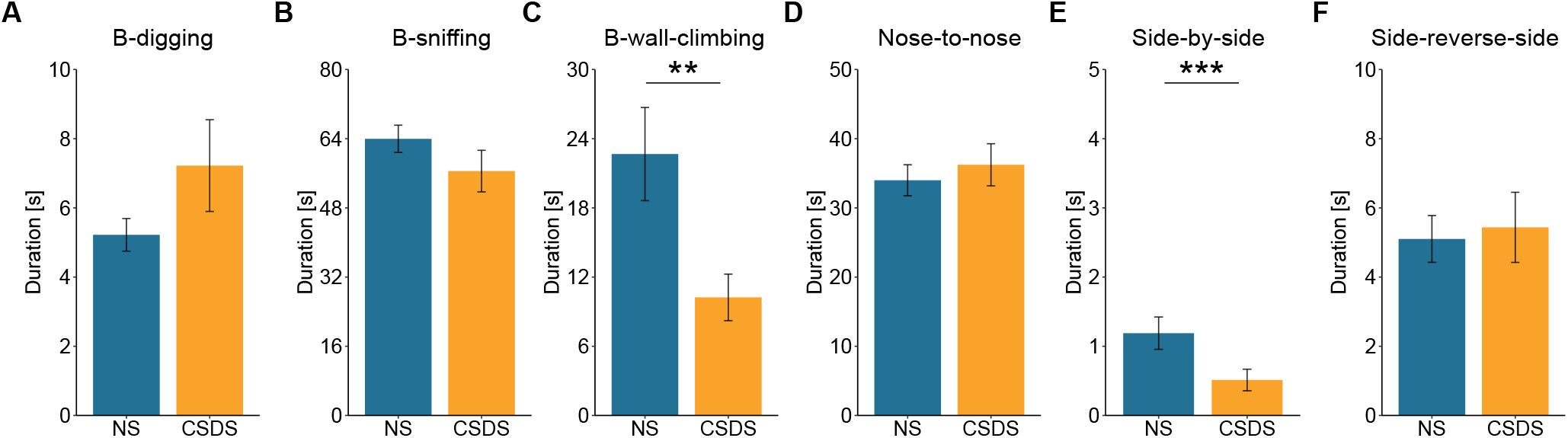
DeepOF other behavioral classifiers in the social interaction task for 10 min total duration. A) B-digging is not significantly different between stressed and non-stressed animals. Wilcoxon test: W=331.5, p=0.735. B) B-sniffing is not significantly different between stressed and non-stressed animals. Welch’s test: T(44.68)=1.30, p=0.201. C) B-wall-climbing is significantly lower in stressed animals. Wilcoxon test: W=496.5, p=0.00988. D) Nose-to-nose is not significantly different between stressed and non-stressed animals. Wilcoxon test: W=334, p=0.769. E) Side-by-side is significantly lower in stressed animals, which only occurred for approximately 1 second in both conditions. Wilcoxon test: W=541.4, p=0.00068. F) Side-reverse-side is not significantly different between stressed and non-stressed animals. Wilcoxon test: W=389.5, p=0.499.

**Fig. S5.**
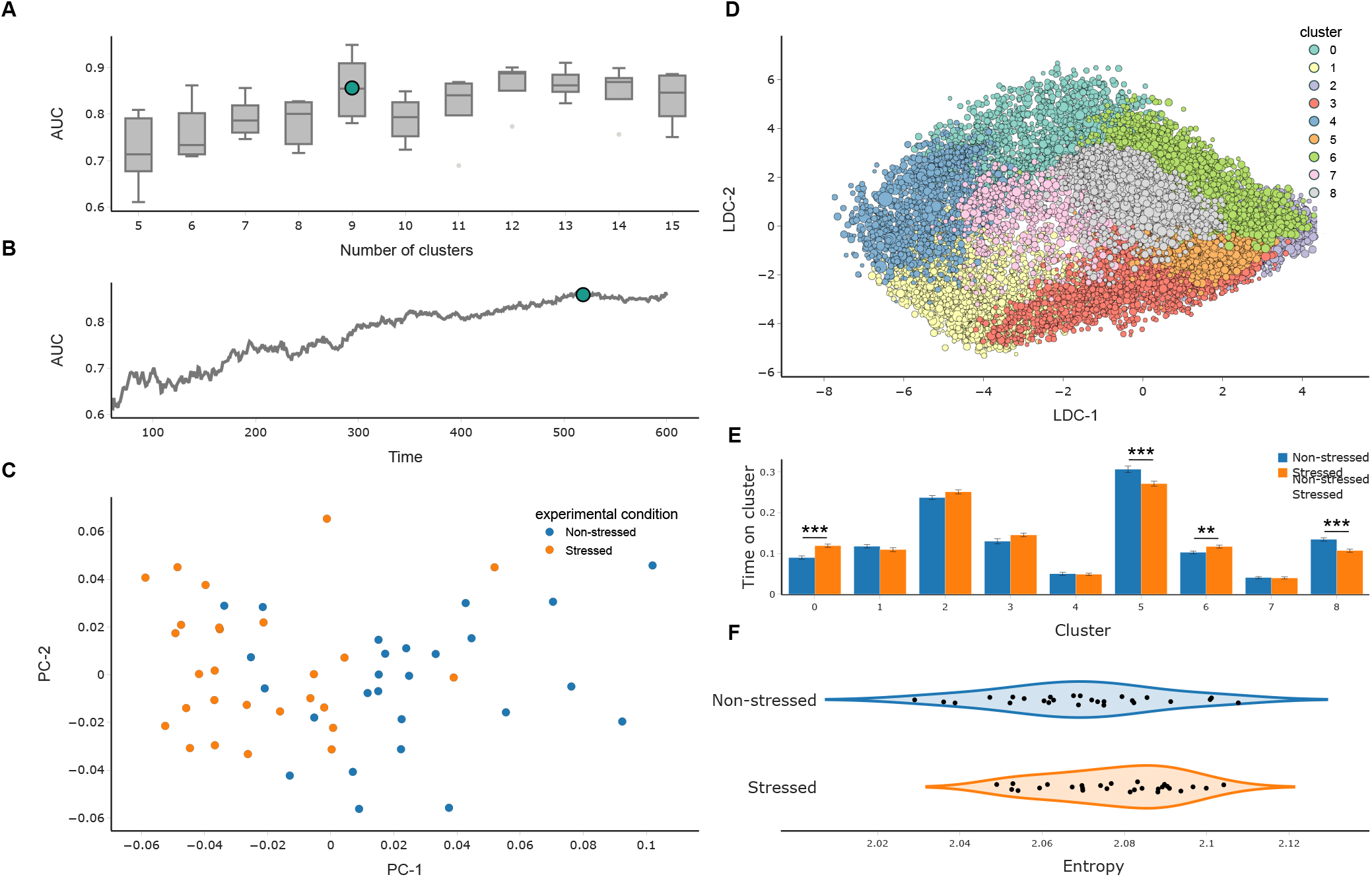
VQVAE unsupervised analyses identify different behavioral patterns between stressed and non-stressed mice during individual open field. A) Cluster selection pipeline results. Models ranging from 5 to 25 clusters were trained in a 10-fold cross-validation loop (only 5 to 15 are shown, for visualization purposes). Discriminability between conditions on the aggregated embeddings (analogous to panel “B)” is reported. The cluster number which maximized this curve (this explaining the most difference between the conditions of interest) was selected. B) Optimal binning of the videos was obtained by measuring the performance of a linear classifier on discriminating between both conditions across a growing window taking a range from the first 60 to 600 seconds for each video (grey curve). Higher values correspond to more distinguishable distributions (i. e., more different behavioral profiles across conditions). We observe a maximum at 8.68 minutes (dark green marker), suggesting that the difference across conditions is sustained across the entire experiment, making binning unnecessary. C) Representation of the aggregated unsupervised embedding for the optimally discriminant bin per experimental video, colored by condition. Each 10-minute-long video was fed to a trained VQVAE model using DeepOF’s unsupervised pipeline. Aggregated embeddings were constructed as the time spent on each defined cluster (yielding a 9-dimensional vector). Dimensionality was further reduced using PCA for visualization purposes. D) Unsupervised embedding of all automatically ruptured video fragments. Different colors correspond to different clusters as recognized by DeepOF. Dimensionality was further reduced using LDA for visualization purposes. E) Cluster population per experimental condition in the first optimal bin (8.68 minutes). Reported statistics correspond to a 2-way Welch t-test corrected for multiple testing using Bonferroni’s method across both clusters and bins. F) Stationary distribution’s entropy of the transition matrices provided by the unsupervised pipeline within DeepOF. No differences were detected between the conditions (t-test p-value=0.08).

**Fig. S6.**
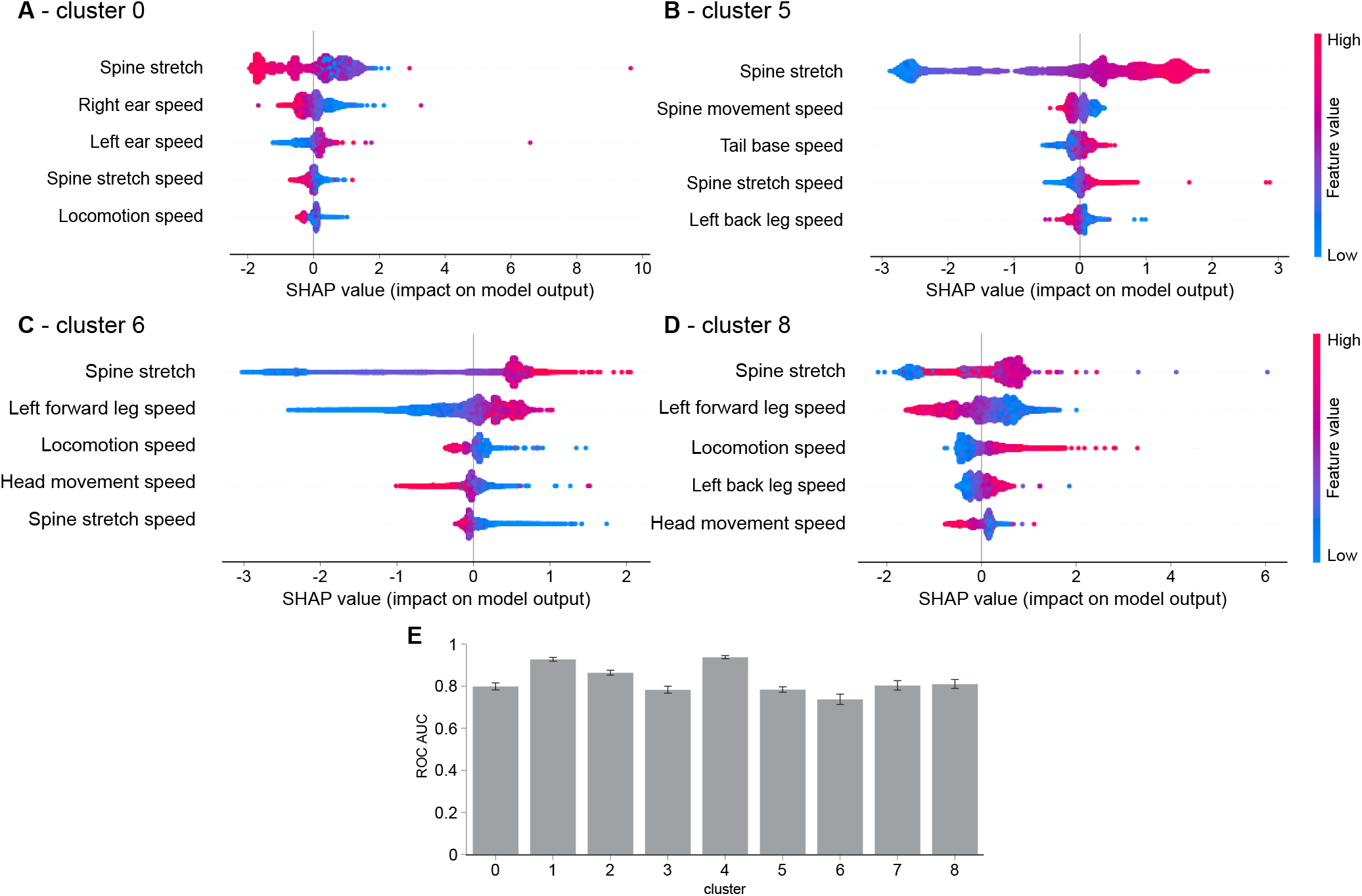
SHAP analysis of unsupervised cluster assignments in the single open field data. A–D) Beeswarm plots for clusters 0, 5, 6, and 8 obtained by the previously discussed social interaction experiments. Gradient boosting machines were trained to map from a predefined set of time series statistics (including locomotion and individual bodypart speeds, distances, and angles) to the previously obtained cluster assignments. The plots depict the first 5 most important features for each classifier, in terms of the mean absolute value of the Shapley additive values (SHAP). F) Performance of the involved gradient boosting classifiers in terms of the area under the ROC curve. Data were standardized, and the minority class was oversampled using the SMOTE algorithm, to correct for class imbalance.

## Notes

https://gitlab.mpcdf.mpg.de/lucasmir/deepof

### Competing Interest Statement

The authors have declared no competing interest.

